# A CRISPR Interference System for Inducible Gene Knockdown in soil bacterium *Sinorhizobium meliloti*

**DOI:** 10.1101/2025.09.23.678109

**Authors:** Francisco J. Guerra-Garcia, Siva Sankari

## Abstract

Symbiotic relationships have an important role in most life forms, but the molecular and cellular processes that establish and maintain these harmonious interactions remain largely unknown. The relationship between leguminous plants and rhizobial bacteria is a classic example of symbiosis, where the bacterium converts atmospheric nitrogen to plant-usable ammonia in exchange for fixed carbon and nutrients. Some legumes such as *Medicago truncatula* has evolved a set of small peptides that exploit this relationship, turning its bacterial partner, *Sinorhizobium meliloti*, into a terminally differentiated bacterium that loses its capability to survive outside the host. However, the mechanisms of how this transformation happens remain elusive due to the absence of high-throughput tools for targeted gene knockdowns in the bacterium. To overcome these limitations in the plant-rhizobia field, we developed an inducible CRISPR-interference knockdown system which can reversibly block the transcription of a target gene through the combined action of a deactivated-Cas9 (dCas9) and single-guide RNAs (sgRNAs). We used a taurine-inducible promoter to achieve fine-tunable expression levels of dCas9 in free-living *S. meliloti* and demonstrated that this tool is suitable for the study of essential genes that could be involved in the symbiotic process, including *hemH, dnaN and ctrA*. Our cost-effective inducible CRISPRi strategy will contribute to understanding the molecular mechanisms underlying legume-rhizobia symbiosis, ultimately allowing soil improvement and reducing chemical fertilizers usage while meeting global food demands.

## Introduction

Symbiotic relationships are found in most life forms, and their mutual benefits often play a central role in the development and survival of both the host and the symbiont. Rhizobia are Gram-negative alphaproteobacteria that can be found in the soil and inside root nodules of leguminous plants establishing a symbiotic association^1^. In this symbiosis, soil bacteria convert atmospheric nitrogen to plant-usable ammonia within the plant cell and receive fixed carbon and other nutrients in exchange. This process, known as biological nitrogen fixation (BNF), helps legumes to overcome the scarce bioavailability of nitrogen in the soil, a crucial element for the survival and growth of plants. This symbiotic relationship not only improves soil fertility but also reduces the need for chemical fertilizers that can pollute ecosystems and contribute to climate change. Despite BNF being a sustainable alternative to meet global food demands, the study of rhizobia is severely limited by the absence of suitable genetic tools.

In recent years, different approaches have been used for genetic modification in rhizobia, with homologous recombination with counter-selectable markers being the most popular^2,3^. For this strategy, 400 to 600 bp long homology arms, complementary to the target region, are designed and cloned into a suicide vector to generate insertions and deletions. However, this approach is time-consuming and limited by the recombination efficiency of the bacteria. Even in the most studied model rhizobium *Sinorhizobium meliloti*, unfortunately, recombination efficiency is only moderate and highly dependent on whether it is targeted to its chromosome or the mega plasmids *PsymA* and *PsymB*^4^. To overcome this limitations, some site-specific recombination methods have been developed in *S. meliloti*^5–7^, but these often demand tedious and single-use experimental designs involving the use of multiple plasmids and yielding disparate outcomes.

For genome-wide studies and high-throughput screenings, the commonly used genetic tool is Tn5 and other transposon-mediated strategies, such as Tn-seq^8^. Nevertheless, Tn5 mutagenesis is known to have its own insertion bias towards long genes and AT-rich regions^9^, which is inconvenient since rhizobia has on average a GC content higher than 60%. Additionally, random and double insertions lead to ambiguous results and complicated analysis.

More recently, CRISPR/Cas systems, such as Cas9^10^, Cas12e^11^ and Cas12k^12^, have been introduced in *Sinorhizobium meliloti* for precise and targeted insertions, deletions and base-resolution editing. Yet, most of this CRISPR/Cas strategies rely on inefficient endogenous machinery, including DNA repair and recombination. Moreover, all the above mentioned tools generate permanent modifications in the genome of free-living rhizobia and require extensive genotyping. Hence, they cannot be used to study essential genes or to selectively disrupt genes when bacteria are associated with the host. Previous studies in *S. meliloti* have reported 832 genes to be essential for growth^8^, making a 13.38% of its coding sequences inaccessible with the current genetic tools because permanent deletions are deleterious or will often result in consequent suppressor mutations that generate misleading phenotypes.

This inability to perturb essential genes and manipulate rhizobia, especially after it has successfully established endosymbiosis, constitutes a major limitation which is detrimental for the study of mechanisms that establish and maintain symbiosis. An example of a symbiotic process which study is restricted by the lack of suitable engineering tools is terminal bacteroid differentiation. Some legumes belonging to the inverted-repeat lacking clade (IRLC), like *Medicago truncatula*, can exploit the symbiotic relationship by secreting a group of small nodule cysteine-rich (NCR) peptides that transform rhizobial bacteria into highly efficient nitrogen-fixing bacteroids^13^. This process, known as terminal bacteroid differentiation (TBD), shapes the bacteroid with unique features, including cell cycle arrest, increased ploidy, increased size and morphology changes. Previous studies have shown that rhizobia undergoing this transformation and acquiring these features have higher nitrogen fixation efficiency than the ones that do not^14^. However, the molecular mechanisms underlying TBD and the action of NCR peptides remain unexplored^15^. This is mainly because of the lack of tools to study essential genes involved in the process.

The goal of this work is to solve most of the limitations mentioned above while keeping a simple, cost effective, fast and efficient design for reversible and selective gene targeting in rhizobia. For that purpose, we have developed a CRISPR-interference (CRISPRi) system in *Sinorhizobium meliloti* for selective gene knockdowns with a taurine inducible promoter. We demonstrate that our taurine inducible CRISPRi approach is suitable for reversible knockdown of essential genes as well as for other useful strategies including dual targeting and operon knockdowns. While toxicity associated to Cas9 expression has been reported in other studies^12^, we show that our CRISPRi system does not display any undesired growth defects. The results and phenotypes observed using this genetic tool resembled previously described outcomes but also led to the discovery of new candidates regulated by essential genes that have not been reported before.

## Results

### Design of a taurine-inducible CRISPRi system in free-living rhizobia

The CRISPR-interference system consists of a deactivated-Cas9 (dCas9) and a single-guide RNA (sgRNA) that contains a 20 base pair long sequence complementary to the target sequence^16^. When targeting the promoter of a gene, the binding of dCas9 to the promoter region does not generate any DNA sequence modification but it is enough to physically block the transcription machinery and therefore inhibit mRNA production. To have full control on the expression of dCas9, we integrated the coding sequence of a dCas9 derived from *Streptococcus thermophilus* (Sth3-dCas9), which has been previously tested in other alphaproteobacteria^17^, in the symbiotic plasmid B (*PsymB*) of *S. meliloti* under the taurine inducible promoter *P*_*tauA*_ in the *tau* locus^18^ (Figure 1a). For gene targeting, sgRNAs were designed by first identifying 20-bp long sequences preceding PAM sequence (NGGNG) instances in a 220-bp window from -200 to +20 of the start codon of the target gene. From these potential sgRNAs, sequences closer to the ATG and targeting the coding strand were prioritized. Chosen sgRNA sequences were checked for possible off-targets in the *S. meliloti* 1021 genome and then cloned upstream of a dCas9 handle and expressed from a high copy plasmid under a constitutive promoter.

**Fig. 1.**
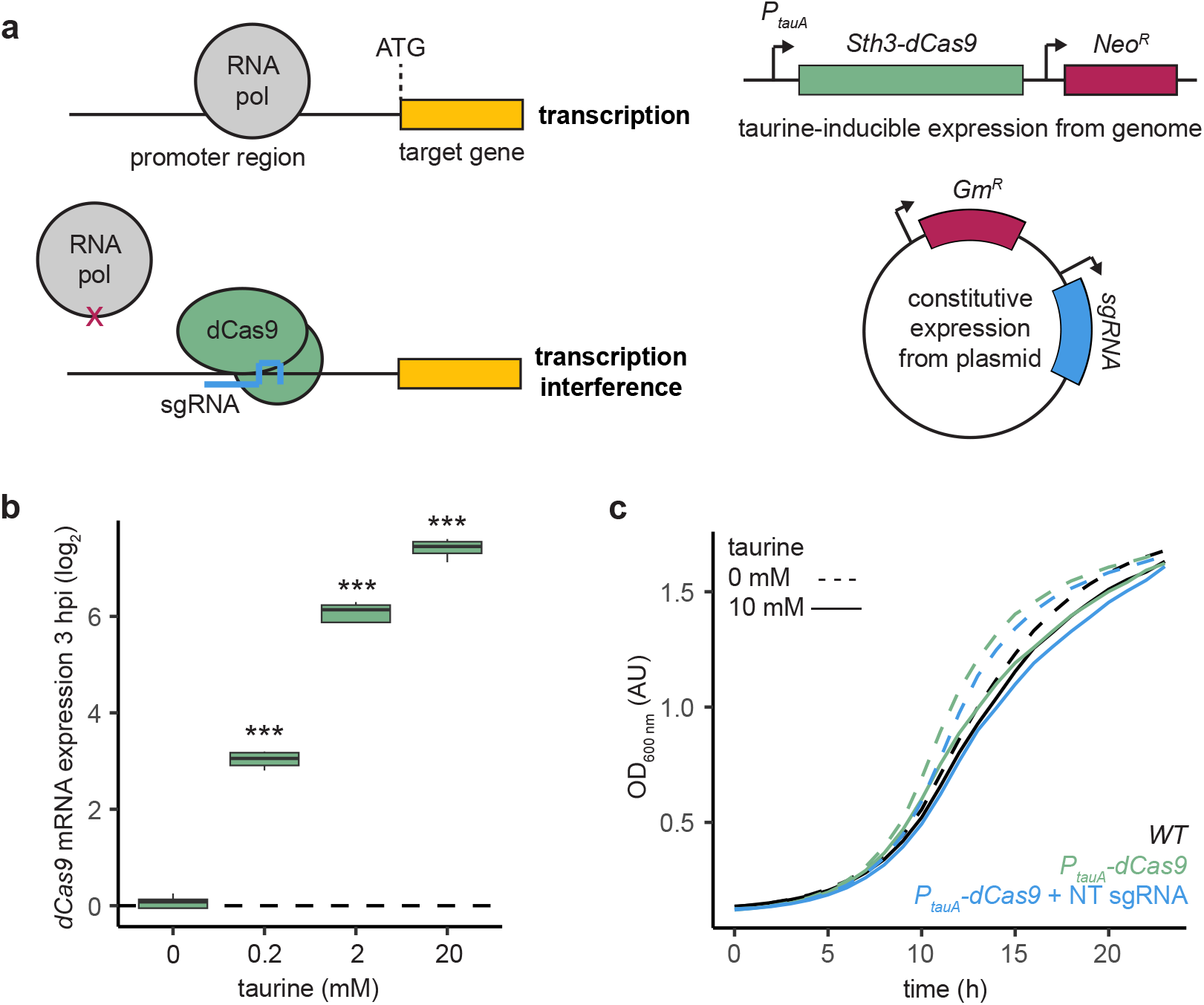
CRISPRi system for inducible gene knockdown in *Sinorhizobium meliloti*. **a**. Representation and design of CRISPR interference system for transcription inhibition. The coding sequence of a deactivated Cas9 derived from *Streptococcus thermophilus* (*Sth3-dCas9*) is integrated in the *S. meliloti* genome at the *tau* locus under the *tauA* promoter for taurine-inducible expression. The single-guide RNA (sgRNA) is expressed constitutively from a plasmid. **b**. *dCas9* mRNA levels measured by RT-qPCR at 3 hours post induction (hpi), normalized to *smc00128* levels. Expression levels with different concentrations of taurine are represented as log_2_ values relative to non-induced control. **c**. Effect of the expression of *Sth3-dCas9* and non-targeting (NT) sgRNA on growth curves of *S. meliloti* 1021, generated with hourly measures of OD_600 nm_.

To test whether dCas9 expression is responsive to taurine induction, we grew *S. meliloti P*_*tauA*_*-dCas9* in media with increasing concentrations of taurine ranging from 0 to 20 mM and measured *dCas9* mRNA relative expression through RT-qPCR. We used the *smc00128* gene for normalization since its expression remains constant in the conditions tested. We show that mRNA levels of *dCas9* were significantly higher in the presence of taurine in the media at 3 hours post-induction (hpi) compared to uninduced samples (Figure 1b). Additionally, we observed increasing expression levels with increasing concentrations of taurine, from an 8-fold change with 0.2 mM taurine to over a 128-fold change with 20 mM taurine. The induction and presence of an HA-tagged dCas9 protein was then confirmed by western blot using an HA antibody (Figure S1a). Increasing protein levels were observed with increasing concentrations of taurine.

Finally, since previous studies have reported that overexpression of the *Streptococcus pyogenes* Cas9 protein can cause cell toxicity and growth inhibition^12^, we monitored the growth of our *S. meliloti P*_*tauA*_*-dCas9* strain and a strain containing both *P*_*tauA*_*-dCas9* and a non-targeting (NT) sgRNA in the presence and absence of 10 mM taurine for 24 hours. No significant changes were observed in the growth rate of the mentioned strains in the tested conditions when compared to wildtype *S. meliloti* (Figure 1c), ruling out negative growth effects associated to dCas9 or sgRNA expression.

### Gene targeting using the inducible CRISPRi system

To test the efficiency of the *P*_*tauA*_*-dCas9* system, we designed an sgRNA complementary to the promoter region of the gene *hemH*. HemH encodes ferrochelatase, an enzyme that catalyzes the ferrous insertion into protoporphyrin IX, which is the last step in the heme synthesis pathway, essential for iron metabolism^19^. Without the addition of external heme, *hemH* knockout mutants are not viable due to the lack of endogenous heme. We first assessed the cell viability of our *hemH*-targeting strain by performing a serial dilution spot assay in plates containing 0 or 10 mM taurine. Compared to the *P*_*tauA*_*-dCas9* strain expressing an NT sgRNA, the viability of the *hemH* sgRNA expressing strain showed a 10^4^-fold decrease only in plates containing taurine (Figure 2a). To check whether the diminished viability was a consequence of targeting *hemH*, we grew the knockdown strain in media with and without taurine for 24 hours and quantified protoporphyrin IX accumulation every hour by measuring red fluorescence (408/635 nm). We observed increased protoporphyrin IX fluorescence only when *hemH* sgRNA was expressed in the presence of taurine (Figure 2b).

**Fig. 2.**
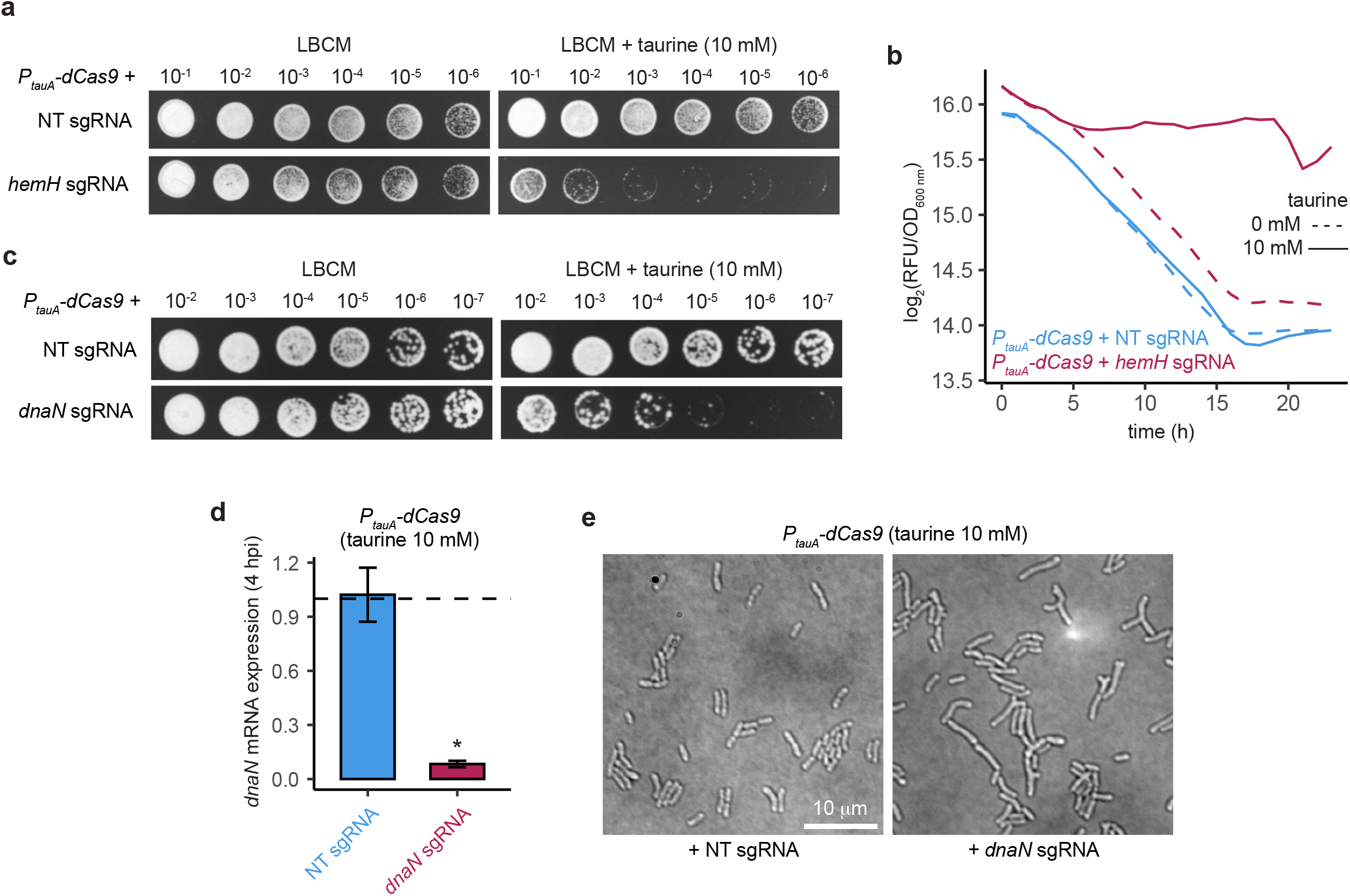
Targeting essential genes with the inducible CRISPRi system. **a**. Viability of strains expressing *P*_*tauA*_*-dCas9* plus NT or *hemH* targeting sgRNAs. Shown are serial dilutions of each strain on LB media supplemented with CaCl_2_ and MgSO_4_ (LBCM) after 3 days at 30°C. **b**. Accumulation of protoporphyrin IX in *hemH* downregulated strains over time. Protoporphyrin IX values are represented as red fluorescence (wavelength 635 nm) intensity normalized by growth (OD_600 nm_) in log_2_ scale. Measures were taken hourly. **c**. Viability of strains expressing *P*_*tauA*_*-dCas9* plus non-targeting (NT) or *dnaN* targeting single-guide RNAs (sgRNAs). Shown are serial dilutions of each *S. meliloti* strain on LB media supplemented with CaCl_2_ and MgSO_4_ (LBCM) after 3 days at 30°C. **d**. *dnaN* knockdown efficiency. mRNA levels were measured by RT-qPCR at 4 hours post induction (hpi), normalized to *smc00128* levels. **e**. Cell branching and elongation observed in *dnaN* knockdown cells using CRISPRi.

To assess the efficiency in a completely different metabolic pathway, we designed a sgRNA targeting *dnaN*, a gene that encodes the *β*-sliding clamp protein, essential for DNA replication^20^. A spot assay showed decreased viability in the *P*_*tauA*_*-dCas9* strain expressing the *dnaN* sgRNA, only in the inducing conditions (Figure 2c). We also detected a significant reduction in mRNA levels of *dnaN* through RT-qPCR analysis (Figure 2d). Additionally, by taking widefield microscopy images, we observed that cells expressing *P*_*tauA*_*-dCas9* + *dnaN sgRNA* looked elongated and were significantly bigger than *P*_*tauA*_*-dCas9* + NT *sgRNA* (Figure 2e). Cell area was segmented and quantified from microscopy images and a significant size increase was confirmed (Figure S1b). This may be due to a reduction in DNA replication while the cell growth continues. Altogether, these results demonstrate that the taurine-inducible dCas9 can be used reliably for creating gene knockdowns in free-living *S. meliloti*.

### Knockdown of the essential cell cycle regulator *ctrA*

CtrA is considered a master regulator of cell cycle and cell division in *S. meliloti* and other alphaproteobacteria^21^. Study of the functionality of CtrA in *S. meliloti* has been hampered due its essentiality, which prevents the generation of stable and conventional deletions or mutations^22,23^. We decided to target *ctrA* to test whether we could resemble the phenotype and features reported by previous studies without the need of creating thermo-sensitive mutants or complementation plasmids. We performed a spot assay to evaluate the viability of a *ctrA*-targeting strain. The results showed that *P*_*tauA*_*-dCas9 S. meliloti* expressing sgRNAs targeting *ctrA* had a 10^4^-fold decrease in viability in plates containing taurine 10 mM compared to a strain expressing NT sgRNA (Figure S1c). We then grew our *ctrA* knockdown strain in liquid media and measured the optical density at 600 nm for 24 hours in 1 hour intervals. We observed that the growth rate of the *ctrA* sgRNA expressing strain was significantly lower than the non-induced and non-targeting controls (Figure 3a). By RT-qPCR, we verified that the mRNA levels of *ctrA* were significantly lower at 4 hpi when expressing *ctrA* sgRNA in the presence of taurine (Figure S2a). Since knocking down genes related to cell cycle and cell division often results in changes in morphology and DNA content, we used widefield microscopy to evaluate morphology, and flow cytometry to measure DNA content^23^. We observed abnormal cell morphology at 24 hpi in our *ctrA*-targeting strain (Figure 3b) compared to the uninduced control. We stained 24 hpi *ctrA* knockdown samples with SYTOX Green to quantify DNA content and compared it to uninduced and non-targeting controls. Compared to these controls, *P*_*tauA*_*-dCas9* rhizobia expressing a *ctrA*-targeting sgRNA showed an accumulation of DNA that reached over the equivalent size of 5 copies of genome, confirming defects in cell cycle and division (Figure 3c), as reported previously with a complementation plasmid strategy^23^. Additionally, cell size data coming from flow cytometry also displayed an increased size in the knockdown strain under induction (Figure S2b).

**Fig. 3.**
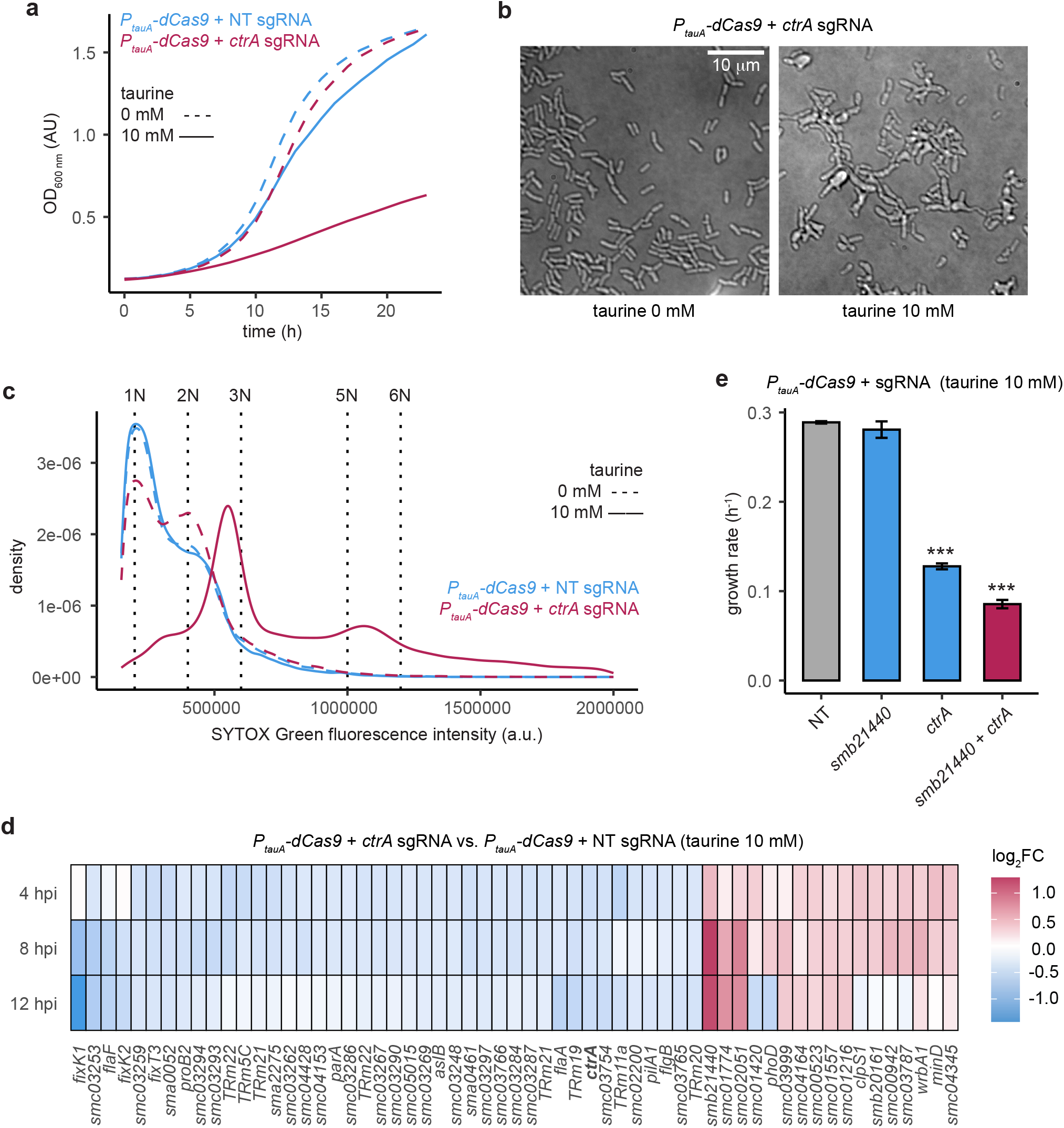
Targeting the cell cycle master regulator ctrA using inducible CRISPRi. **a**. Inhibitory effect on growth in strains expressing *P*_*tauA*_*-dCas9* plus *ctrA* targeting sgRNA, generated with hourly measures of OD_600 nm_. **b**. Morphology defects of *P*_*tauA*_*-dCas9* + *ctrA* sgRNA cells in presence of the inducer versus uninduced cells. **c**. Flow cytometry profiles showing the abnormal DNA content of *ctrA* knockdown cells at 24 hours after induction of the CRISPRi system. DNA content was measure with SYTOX Green dye. The intensity values associated to number of genome copies (ploidy number, N) are represented with dashed black vertical lines. **d**. List of genes significantly down- and up-regulated in at least two out of three time points taken at 4, 8 and 12 hours after induction of *ctrA* knockdown. Expression values are shown as log_2_ fold-change (log_2_FC) of transcripts per million (TPM) in the *P*_*tauA*_*-dCas9* strain expressing *ctrA* sgRNA versus NT sgRNA. Thresholds: FDR < 0.05, |log_2_FC| > 0.4. **e**. Synergistic inhibitory effect on growth rate due to simultaneous knockdown of *ctrA* and *smb21440* with dual targeting, compared to single targeting effects. Growth rate (number of doublings per hour) was calculated from the exponential growth phase of induced *P*_*tauA*_*-dCas9* cells expressing NT, *smb21440, ctrA* or *smb21440* plus *ctrA* sgRNAs.

### Transcriptomic analysis on *ctrA* knockdown rhizobia

While the phenotype of our *ctrA* knockdown strain resembles the one described in previous studies with conditional depletion^23^, we wondered if the transcriptome variations were also related. To address this, we collected *ctrA* knockdown samples at 4, 8 and 12 hpi as well as non-targeting controls at the same time points and performed an RNA-seq analysis. As expected, *ctrA* appeared significantly down-regulated in the three time points, together with over a hundred up- and down-regulated genes (Figure S2c). To narrow down the number of relevant candidates, we made a list with genes that were significantly regulated in at least two of the three time-points. Within this list we identified genes that were previously reported to be regulated by *ctrA*, such as *flaA, flgB* or *pilA1*, as well as other novel genes not reported before (Figure 3d). For validation, we picked five of the strongest regulated genes based on fold change in expression, as well as the previously reported genes *flaA* and *flgB*, and performed RT-qPCR analysis at 4 hpi. The mRNA levels of the tested genes followed the behavior observed in the RNA-sequencing experiment (Figure S2a). Thus, the inducible CRISPRi system allowed the targeting and knockdown of an essential gene and revealed candidate genes that might be regulated by *ctrA* and that have not been described in similar studies performed with different genetic tools. As an example, we highlight the uncharacterized gene *smb21440*, which appeared to be significantly up-regulated in all our experiments when *ctrA* was knocked down, and has not been reported as part of the *ctrA* regulon before. This suggests that *smb21440* may play an important role in CtrA regulon. To test this, we designed an sgRNA targeting the *smb21440* gene promoter and assessed growth under induction conditions. We observed that while the knockdown of *smb21440* alone did not show any effect on growth, the simultaneous knockdown of *smb21440* and *ctrA* together showed a significant decrease of growth rate compared to the knockdown of *ctrA* alone (Figure 3e). This synergistic effect between *ctrA* and *smb21440* suggests that *smb21440* up-regulation is more likely associated to *ctrA* knockdown and not to a nonspecific response to stress.

### Dual targeting with CRISPRi

In bacterial genetic studies, often there is a need to target multiple genes simultaneously to study synergistic effects. To deepen our characterization of the targeting behavior of our inducible CRISPRi system in free-living rhizobia, we tested whether two genes could be targeted simultaneously by co-expressing two different sgRNAs from the same plasmid. Since *hemH* knockdown cells displayed red fluorescence and regular morphology, and *ctrA* knockdown cells showed defective growth and abnormal morphology, we decided to target these two genes with easily assessable phenotypes. We monitored the growth of *S. meliloti P*_*tauA*_*-dCas9* strains expressing *hemH, ctrA, hemH* + *ctrA* or non-targeting sgRNAs in media with and without taurine. We found that only *ctrA* sgRNA *and hemH* + *ctrA* sgRNA expressing cells showed defects in growth in the presence of the inducer (Figure 4a). We confirmed the morphology defects in *ctrA* sgRNA *and hemH* + *ctrA* sgRNA expressing cells through widefield microscopy (Figure 4b). On the other hand, only *hemH* sgRNA *and hemH* + *ctrA* sgRNAs showed protoporphyrin IX accumulation measured through red fluorescence (Figure 4c). This data supports that the inducible CRISPRi system is suitable for targeting multiple genes simultaneously without phenotypic interferences.

**Fig. 4.**
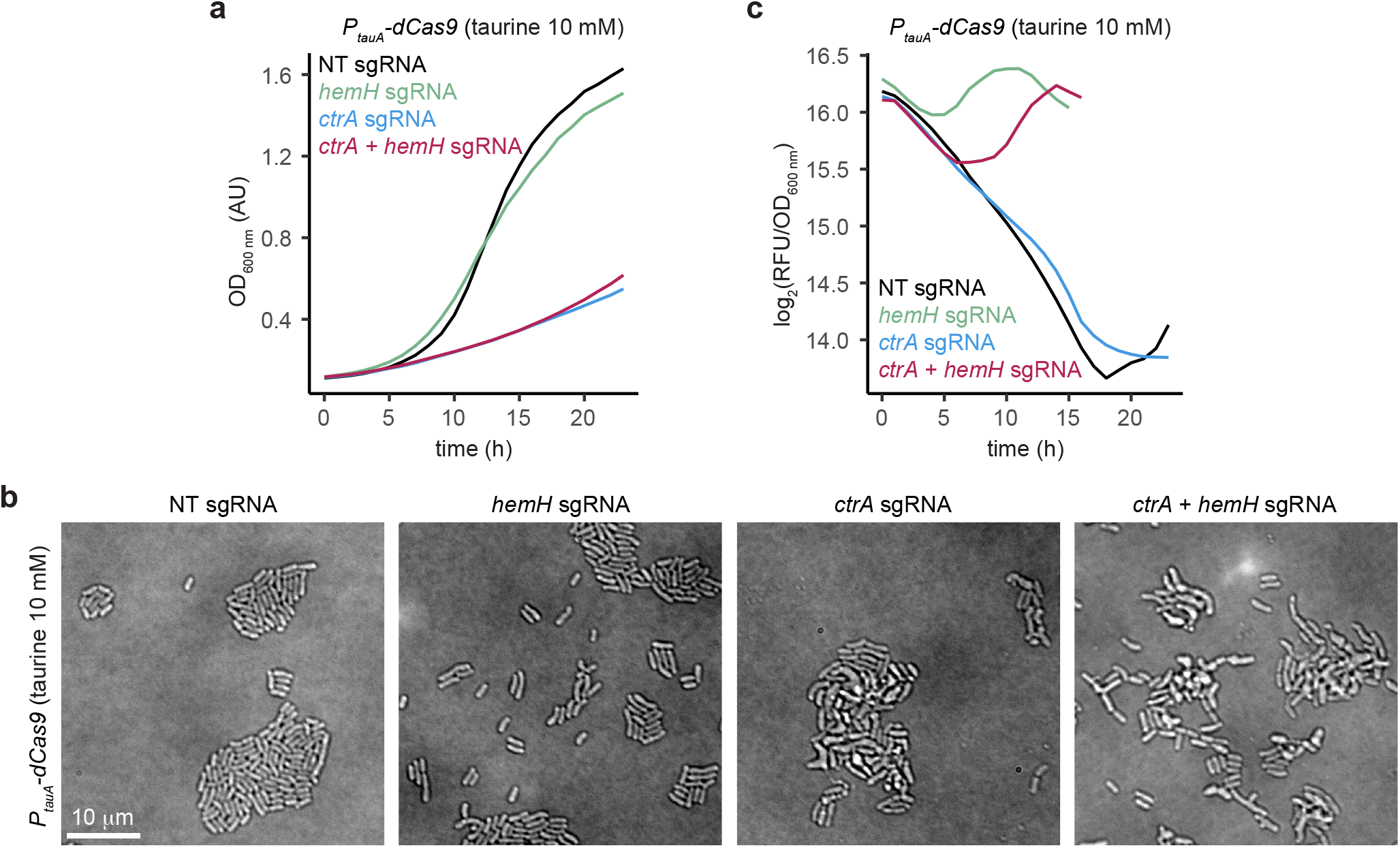
Dual targeting with CRISPRi in *S. meliloti*. **a**. Effect of dual targeting of *ctrA* and *hemH* on growth with *P*_*tauA*_*-dCas9* compared to single targeting with *ctrA, hemH* or non-targeting (NT) single-guide RNAs (sgRNAs), generated with hourly measures of OD_600 nm_. **b**. Accumulation of protoporphyrin IX over time in cells with dual targeting of *ctrA* and *hemH* using CRISPRi. Protoporphyrin IX values are represented as red fluorescence (wavelength 635 nm) intensity normalized by growth (OD_600 nm_) in log2 scale. Measures were taken hourly. **c**. Morphology defects of dual targeting *P*_*tauA*_*-dCas9* cells expressing *ctrA* and *hemH* sgRNAs.

### Reversibility of the inducible CRISPRi system

One of the major strengths of the current design of the CRISPRi system is the inducibility, which allows activation of gene targeting upon addition of taurine to the media. This induced and partial transcriptional knockdown enables the study of essential genes whose knockout would be lethal otherwise. Taking advantage of these features, we wondered whether an induced knockdown of an essential gene could be switched off, and normal function restored upon removal of the inducer. We first evaluated the ability of dCas9 expression to be turned off after removal of taurine. We measured *dCas9* mRNA expression at different time points after adding taurine 10 mM to the media and after removing it with media washes. Interestingly, we observed a significant 24-fold increase in dCas9 expression as soon as 15 minutes post induction (Figure 5a), demonstrating how rapid induction can be achieved with this system in free-living rhizobia. Moreover, washing the taurine from the media at 3 hours post induction resulted in a significant 32-fold decrease in dCas9 expression in 2 hours. After confirming that expression levels of dCas9 can be turned on and off, we assessed the reversibility of a knockdown by designing a recovery experiment. Here we induced *hemH* knockdown with taurine for 6 hours. After that, we washed the cultures and split them into two conditions: taurine-removed (0 mM) and taurine re-added (1 mM). Protoporphyrin IX fluorescence was measured and normalized by comparing the *hemH*-targeting strain and the non-targeting strain at 0, 12, and 24 hours post wash (hpw). We observed significant differences beginning at 12 hpw and, by 24 hpw, the cultures that had the taurine removed from the media displayed more than a 2-fold decrease in fluorescence compared to the ones that still contained taurine (Figure 5b). This suggests that HemH function can be recovered after we wash the taurine, indicating the knockdown system is reversible.

**Fig. 5.**
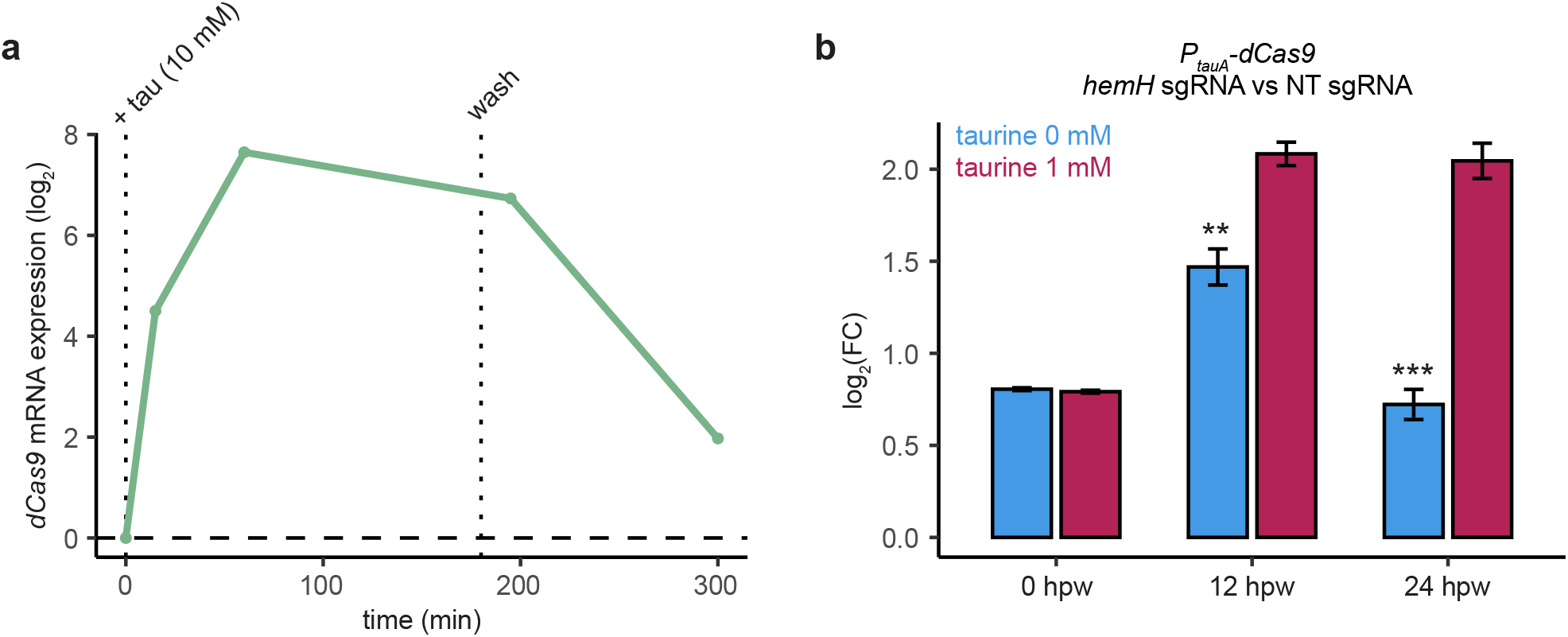
Reversibility and phenotype recovery with the inducible CRISPRi system. **a**. *dCas9* mRNA levels measured by RT-qPCR, normalized to *smc00128* levels. Expression levels at different time points (15, 60, 195 and 300 minutes post induction) are represented as log_2_ values relative to time zero. Inducer was washed off the media at 180 minutes post induction. **b**. *hemH* function recovery after washing the inducer off the media. Recovery is measured as a log_2_ fold change (log_2_FC) of protoporphyrin IX accumulation in *P*_*tauA*_*-dCas9* cells expressing *hemH* sgRNA versus NT sgRNA. Protoporphyrin IX accumulation is calculated as red fluorescence (wavelength 635 nm) intensity normalized by growth (OD_600 nm_). The inducer was washed from the media after 6 hour of induction and recovery was calculated at 0, 12 and 24 hours post wash (hpw) in samples where taurine was added back to the media, and in samples that remained free of inducer.

### sgRNA targeting efficiency and operon targeting with CRISPRi

Our inducible CRISPR-interference system solves most of the limitations that current gene engineering tools display in the rhizobia field. Nevertheless, CRISPRi has some limitations. The efficiency of the target gene knockdown is often affected by several sgRNA features such as relative binding position and targeted strand^12,24,25^. Previous studies have shown that sgRNAs targeting the coding (non-template) strand and sgRNAs targeting promoter regions, especially the predicted RNA polymerase binding site, often yield stronger interference than those targeting the template strand or the coding sequence region. To understand targeting efficiency changes due to sgRNA variability, we tested four sgRNAs targeting different regions and strands of the *hupB* gene promoter, individually and in combinations (Figure 6a). The *hupB* gene encodes a histone-like protein that has been proven to regulate *S. meliloti* cell division through lysine acetylation^26^. By measuring *hupB* mRNA levels with RT-qPCR, we found that two out of the four individual sgRNAs and combinations containing any of them significantly reduced *hupB* expression (Figure 6b). The sgRNAs with higher knockdown efficiency were those that target regions closer to the *hupB* start codon (*hupB* sgRNA 1 and 4). Specifically, the *hupB* sgRNA 1, which yielded the highest interference, targets the coding strand. Another limitation of CRISPRi is that genes within operons cannot be targeted since transcription inhibition is based on promoter targeting. Operons are clusters of genes found in prokaryotes that are transcribed together under the same promoter into a single polycistronic mRNA. While genes within operons cannot be targeted individually, all the genes belonging to the same operon could be potentially knocked down simultaneously by targeting the promoter of the operon. To verify this, we designed an sgRNA targeting the promoter of the operon that contains the genes *nifA, nifB* and *fdxN*. We performed a growth assay of the *P*_*tauA*_*-dCas9* strain expressing a *nifA* sgRNA by measuring optical density every hour for 24 hours in media with and without taurine and compared it to the strain expressing a NT sgRNA. No significant differences were found in the growth rate of the mentioned strains in the conditions tested. This proved that targeting *nifA*, a gene essential for endosymbiotic bacteroids but not for free-living bacteria, does not generate an unexpected growth phenotype (Figure 6c). Then, we measured mRNA levels of *nifA, nifB* and *nifT*, a gene that is immediately downstream of the described operon but that is not part of it. At 6 hpi, in the presence of taurine, we found a significant decrease in mRNA levels of *nifA* and *nifB*, and a significant increase in mRNA levels of *nifT* (Figure 6d). This shows that the inducible CRISPRi system can be used for targeting multiple genes that are transcribed under the same promoter. Additionally, since operons are mainly predicted *in silico* but not tested *in vivo*^27^, this knockdown approach could be useful for validating operons and confirming the genes included in them.

**Fig. 6.**
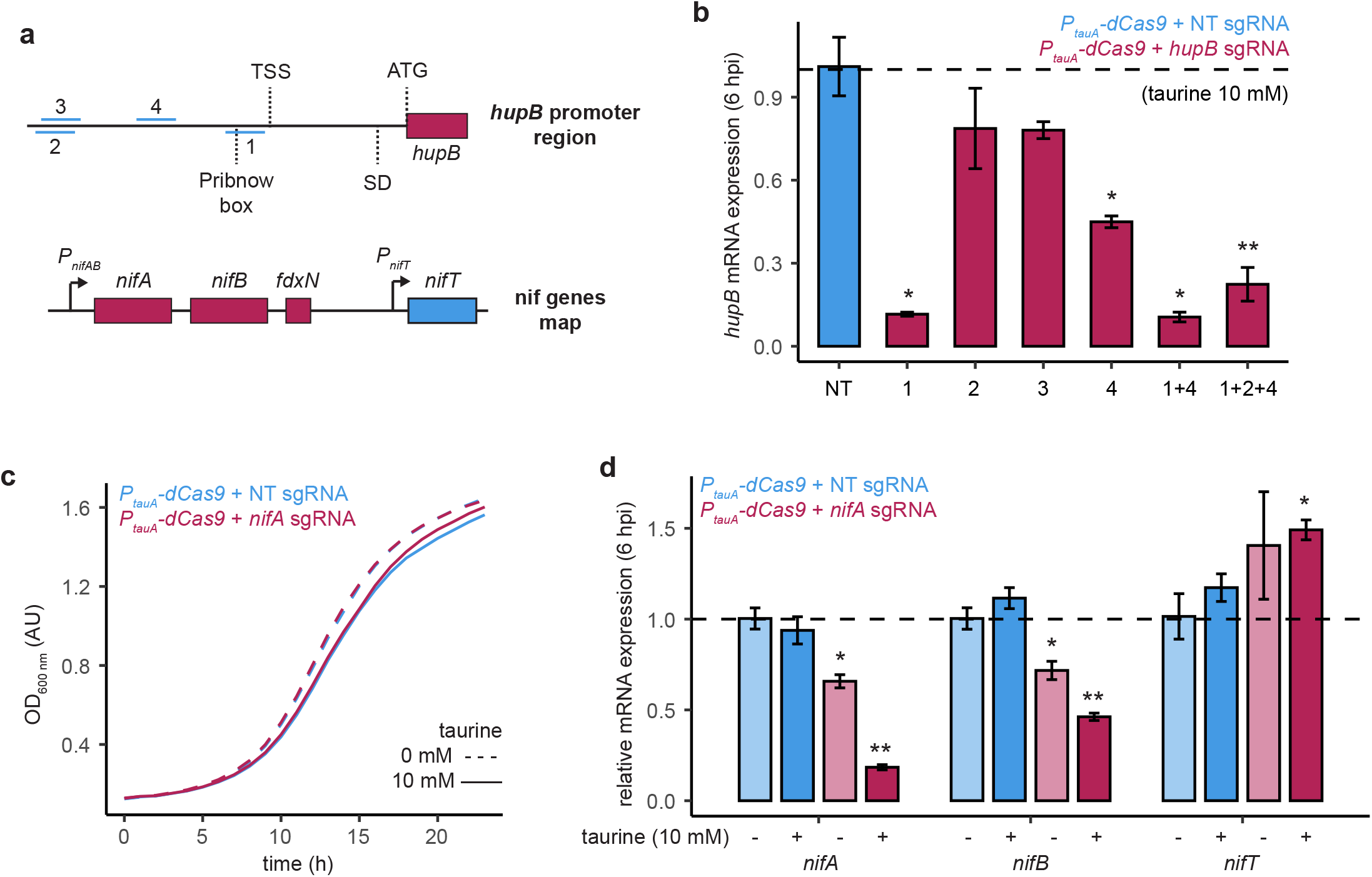
sgRNA targeting efficiency and operon targeting with CRISPRi. **a**. Representation of *hupB* promoter and *nif* genes genomic regions. In the *hupB* promoter region, positions of sgRNAs (numbered from 1 to 4 and represented with blue lines) are shown relative to predicted promoter elements: Pribnow box, transcriptional start site (TSS) and Shine-Dalgarno (SD). In the *nif* genes map, genes expressed under the same promoter are displayed with the same color. **b**. *hupB* knockdown efficiency expressing 4 different *hupB* targeting sgRNAs alone or in combinations. mRNA levels were measured by RT-qPCR at 6 hpi, normalized to *smc00128* levels. **c**. Effect of the expression of nifA targeting sgRNA on growth of PtauA-dCas9 cells, generated with hourly measures of OD_600 nm_. **d**. *nifA, nifB* and *nifT* mRNA levels measured by RT-qPCR at 6 hours post induction (6 hpi), normalized to *smc00128* levels. While *nifA* and *nifB* genes belong to the same operon, *nifT* gene is found downstream of that operon and does not belong to it.

## Discussion

The study of plant-bacteria symbiosis is severely hampered by the absence of suitable genetic tools that allow manipulation of rhizobia at different levels of complexity. We considered most of the limitations that current gene engineering tools have in rhizobia^4^ and introduced a CRISPR-interference system in the model bacterium *Sinorhizobium meliloti*. The deactivated-Cas9 effector derived from *S. thermophilus*, allows targeted inhibition of transcription without generating permanent scars in the genome. To avoid lingering dCas9 expression and achieve time-controlled knockdowns, we made the system fine-tunable by integrating a single copy of the dCas9 coding sequence in the genome under a taurine-inducible promoter. Previous literature has shown that this promoter offers tight control on expression with minimal leaky activity in a wide range of divergent members of alphaproteobacteria, such as *S. meliloti, Caulobacter crescentus* and *Zymomonas mobilis*^18^, suggesting that our CRISPRi system can be easily transferred to most related bacteria. In our hands, the *P*_*tauA*_ promoter in *S. meliloti* yielded dCas9 expression levels that correlated with taurine concentrations and a significant induction effect could be observed as soon as 15 minutes after the addition of the inducer.

To make our inducible CRISPRi approach more accessible and cost-effective, we used a high-copy plasmid that allows the cloning of new single-guide RNAs with a one-step method that involves ordering one single primer. Through the combined action of the inducible *P*_*tauA*_*-dCas9* and targeting sgRNAs we showed that this system is suitable for the knockdown of essential genes in *S. meliloti* such as *hemH, dnaN* and *ctrA*.

Despite the efficiency and versatility of CRISPRi tools, the use of CRISPR/Cas systems often comes with limitations and concerns. We have sought to address these concerns in our tool set in the following ways. i) Previous studies have demonstrated that the expression of a dCas9 derived from *S. pyogenes* can cause toxicity and growth inhibition^12^. We show that the expression of the *S. thermophilus*-derived dCas9 alone or in combination with a non-targeting sgRNA did not cause any detrimental effects in *S. meliloti* growth. ii) Some studies suggest that when targeting sgRNAs, stalling of the replisome due to collision with CRISPR-dCas9 bound to target DNA could occur^28^. However, we demonstrate that targeting genes like *nifA*, which is non-essential for free-living rhizobia, would yield a significant knockdown while keeping cell growth, and therefore replication, unaffected. It is possible that replication interference may be more problematic in fast-growing bacteria and when using sgRNAs targeting genes within regions of replication origin. iii) Other major concerns include off-target effects (undesired transcription interference on genes other than the target) and leaky activity (transcription interference on the target in non-induced conditions). Since the level of restriction of the PAM sequence from *S. thermophilus* CRISRP-Cas system (NGGNG) is higher than other commonly used systems, e.g. *S. pyogenes* CRISRP-Cas9 (NGG), and considering that each sgRNA sequence was checked for complementarity with the whole *S. meliloti* genome, off-target activity was not expected and was not observed to the best of our knowledge. Regarding leaky expression of dCas9 from the *P*_*tauA*_ promoter, we have detected mRNA decrease of the target gene in non-inducing conditions to some extent. Nevertheless, this effect was often result of specific sgRNAs and the associated phenotypes were only displayed in the presence of the inducer. We attribute this phenomenon to the combination of minimal leaky expression of dCas9 and a highly efficient targeting sgRNA. We addressed this issue by testing multiple sgRNAs against the target genes and choosing those that yielded a strong knockdown only when taurine is added to the media.

We also showed that multiple genes can be targeted simultaneously, allowing the study of genes with complementary functions, interacting genes, metabolic pathways, etc. Similarly, multiple sgRNAs against the same target gene can be co-expressed when their efficiency is unknown. This approach, evident from targeting the gene *hupB*, can be very informative and yield significant knockdowns without the need of testing them individually. On the other hand, while genes within operons cannot be individually targeted because they lack a specific promoter, we have shown with the *nifA* operon that our CRISPRi strategy can be successfully used to simultaneously knockdown all the genes belonging to the same operon by targeting its promoter. However, we acknowledge that for targeting specific gene in an operon, we have to resolve to other methods of gene disruption.

Unlike traditional gene deletions and knockouts, the proposed CRISPRi system is particularly well-suited for knockdowns of essential genes. The study of essential genes like *ctrA*, a cell cycle master regulator^21^, has been hampered by the difficulty of generating viable and stable mutant strains. Other research groups have achieved conditional mutations by using a *ctrA* thermo-sensitive allele or a conditional *ctrA* depletion strain^22,23^. While these strategies can succeed, the design is complicated and must be developed on a case by case basis. We are able to recapitulate most of the known features of CtrA mutants by targeting this gene with the inducible CRISPRi system in *S. meliloti*. By using this method and performing transcriptomic analysis on *ctrA* knockdown cells, we reported new potentially *ctrA*-regulated genes such as *smb21440*, which showed an interesting synergistic growth defect. Additionally, we demonstrated that knockdown of essential genes can be time-controlled and reversible. After inducing transcription interference of *hemH* and observing its associated phenotype, removal of the inducer from the media resulted in reversion of the phenotype and HemH function was restored.

To complete the CRISPRi system and fill the remaining gaps in rhizobial genetic manipulation, our future work plans to leverage this tool by adding two new functionalities: i) create and establish a genome-wide sgRNA library suitable for high-throughput screenings in free-living *S. meliloti*, ideal for finding candidate genes involved in a given selection condition; and ii) use symbiosis-inducible promoters^29^ for the expression of the CRISPRi system exclusively in bacteroids after rhizobial infection in plant roots, achieving gene targeting at different stages of symbiosis. Altogether, the inducible CRISPRi strategy developed in this work proved to be valuable as a versatile gene engineering tool. We strongly believe that this tool will be a valuable contribution to the field and open opportunities for deeper examination of the intricacies of the molecular mechanisms underlying soil bacteria biology and legume-rhizobia symbiosis.

## Materials and methods

### Design of CRISPRi sgRNAs

Single-guide RNAs (sgRNAs) were designed for each target promoter with the following considerations. All the PAM sequence (NGGNG) instances in the promoter (220-bp window from -200 to +20 of the start codons of the annotated coding region) were identified, and 20 bp were extracted from the ending 5’ of the PAM sequence as potential targeting sequences. Any sgRNA with potential off-targets effects in the *S. meliloti* 1021 genome were excluded. sgRNAs targeting the coding strand were prioritized over template strand-targeting sgRNAs. All the sgRNA sequences used in this study can be found in Table S1.

### Plasmid construction

All the primers and plasmids used in this study are listed in Table S2 and Table S3. For constitutive expression of sgRNAs under a *P*_*constitutive*_, chosen individual sgRNAs were cloned into the plasmid fgp004, a vector derived from Addgene #133339^17^ by switching Neo^R^/Kan^R^ to Gen^R^ (fgo039-fgo040, fgo041-fgo042). The sgRNA cloning consisted in a divergent PCR with a common reverse oligo (fgo004) and a variable forward oligo that contained the chosen 20 bp sgRNA sequence and a common 22 bp sequence. For dual targeting, a plasmid containing two sgRNA cassettes was generated from individual sgRNA plasmids with primers fgo065-fgo066 and fgo067-fgo068. The PCR product was then phosphorylated, circularized and transformed in chemically competent *E. coli* DH5-alpha. For integration of Sth3-dCas9 into the *S. meliloti tauA* locus, the HA-tagged *P*_*tauA*_*-Sth3-dcas9* cassette was cloned in a pUC119-derived suicide vector (Neo^R^/Kan^R^ Amp^R^) to generate fgp005.

### Strain construction

The 1021 strain of *S. meliloti* was used as the wild-type strain background for all strain constructions in this study, which can be found in Table S4. The insertion of Sth3-dcas9 (from *S. thermophilus*) into the *Sinorhizobium* genome was done via homologous recombination at the *tauA* locus under the *tauA* promoter. The dCas9-containing plasmid and the sgRNA-expressing vectors were introduced in *S. meliloti* through conjugation following a triparental mating protocol. For mating, saturated primary cultures of donor *E. coli* DH5-alpha, helper *E. coli* HB101-pRK600 and acceptor *S. meliloti* were spun down and washed with LB media to remove selecting antibiotics. The three cultures were then suspended in different volumes of LB media to reach similar optical densities. Next, 120 *μ*l of a 1:1:1 strain mix was spotted on LBCM agar plates and incubated at 30 °C overnight. The mating spots were then streaked on selection plates supplemented with the appropriate antibiotics and incubated at 30 °C for 3 days. Single colonies were picked and re-streaked again in selection plates. The resulting clones were tested with colony PCR using specific primers to check for integration in the expected genomic locus (fgo002-fgo025), or for plasmid presence (fgo004-fgo039).

### Bacterial culture conditions

*Sinorhizobium meliloti* strains were grown in lysogeny broth (LB) supplemented with 2.5 mM CaCl_2_ and 2.5 mM MgSO_4_ (LBCM) at 30 °C and 250 rpm horizontal shaking unless otherwise noted. Primary cultures from glycerol stocks were grown in 5 ml LBCM with the appropriate antibiotics until they reached saturation (30 to 48 h). Secondary cultures were grown in LB overnight from primary cultures until they reached exponential phase (0.2 to 0.6 OD_600 nm_). *Escherichia coli* strains were grown in LB with the appropriate antibiotics at 37 °C and 250 rpm horizontal shaking for 16 h. The following antibiotics were used at the indicated concentrations: streptomycin (Strep), 200 *μ*g/ml; neomycin (Neo), 100 *μ*g/ml; gentamicin (Gent), 60 *μ*g/ml; tetracycline (Tet), 10 *μ*g/ml; kanamycin (Kan), 25 *μ*g/ml; chloramphenicol (Chl), 50 *μ*g/ml; ampicillin (Amp), 100 *μ*g/ml. Unless otherwise noted, induction of dCas9 expression was performed by adding 10 mM taurine.

### Spot assays

Secondary cultures of *S. meliloti* at exponential phase were 10-fold serially diluted. 10 *μ*l of each dilution was spotted onto LBCM plates with or without taurine and incubated at 30 °C for 3 days. The plates were then imaged with an Interscience Scan 500 colony counter.

### Growth assays and fluorescence measurement

All growth and fluorescence curve experiments were performed in a Tecan Spark plate reader using flat-bottom 96-well plates. In each well, 10 *μ*l of saturated *S. meliloti* primary culture was added to 190 *μ*l of growth media with inducer present if applicable. The plates were programmed to continuously shake at 180 rpm, and the temperature was maintained at 30 °C. For growth, LB media was used, and optical density was measured at 600 nm every 60 min for 24 hours. For protoporphyrin IX, glucose-salts-yeast extract (GSY) media was used to minimize background fluorescence, and fluorescence was measured using 408/635 nm excitation/emission wavelengths with a bandwidth of 20 nm and manual gain every 60 min for 24 hours.

### Flow cytometry

From the 96-well plates used for growth curve assays, after the 24 hours of incubation, 4 replicates of 200 *μ*l were merged. 100 *μ*l of the replicate mix was fixed by adding 900 *μ*l of 100% ethanol. Fixed samples were spun down for 5 min at 6500 rpm and suspended in 1 ml of 50 mM citrate buffer containing RNase A. Samples were incubated at 50 °C for 2 hours. 190 *μ*l of sample was mixed with 10 *μ*l of 75 *μ*M SYTOX Green and incubated at room temperature for 40 min in the dark for DNA staining. Stained samples were run in the Cytek Aurora, and the resulting data was gated using FCS Express 7 and analyzed using a customized script in R 4.4.1.

### Microscopy

For morphology checks, 6 *μ*l of bacterial culture was spotted on agar plugs and let dry. The agar plugs were placed on 8-well chamber slides with the bacteria facing down. Transmitted light (Köhler illumination) microscopy was performed using a Nikon Ti2-E widefield microscope with 1.5x magnifier and a Nikon Plan Apo Lambda 100x/1.45 oil objective. Images were taken with a Photometrics Prime 95B sCMOS camera. Images brightness and contrast were adjusted using Fiji ImageJ. Cell area measurements were taken using a customized algorithm for cellular segmentation.

### RNA isolation and RT-qPCR

Secondary cultures of *S. meliloti* at exponential phase were spun down and suspended in LB with or without taurine and incubated at 30 °C and 250 rpm horizontal shaking for given time intervals. 1.5 ml samples were collected and spun down. Cell pellets were suspended in 140 *μ*l of lysis buffer provided in the MagMAX”-96 Total RNA Isolation Kit (Thermo Fisher Scientific). After bead-beating the lysate, the protocol of the mentioned kit was followed to obtain total RNA. cDNA was synthesized through reverse transcription using SuperScript” IV VILO” Master Mix with ezDNase” Enzyme (Thermo Fisher Scientific). RT-qPCR primer pairs were validated with standard curves generated with serial dilutions of cDNA. The optimal cDNA dilution was chosen according to the detection range of the QuantStudio” 7 Pro Real-Time PCR System. Relative gene expression values were obtained through delta-delta C_q_ method using *smc00128* gene quantification cycle (C_q_) values for normalization since its expression levels did not change in the conditions tested. Normalized values were then expressed as an average of triplicates, with standard deviation represented by the error bars.

### Total RNA-sequencing

Total RNA was isolated from samples following the method described in the RNA isolation and RT-qPCR section. The RNA quality was assessed using Agilent 2100 Bioanalyzer prior to library preparation for RNA-sequencing. RNA libraries were prepared using FastSelect Watchmaker RNA. The libraries were sequenced using G4 from Singular Genomics with a F3 flow cell, for 50 bp long paired reads. Reads were aligned to UCSC *S. meliloti* 1021 genome (gca_000006965) with STAR aligner (version 2.7.10b). Transcripts per million (TPM) values were generated using RSEM (version v1.3.1).

### Immunoblotting

1.5 ml of secondary cultures of *S. meliloti* at exponential phase were spun down and suspended in a mix of 10 *μ*l beta-mercaptoethanol and 90 *μ*l 4x Laemmli sample buffer (Bio-Rad). The sample was then boiled at 95 °C for 5 min and loaded on a 4-15% Mini-PROTEAN TGX Stain-Free™ protein gel (Bio-Rad) for electrophoresis. The electrophoretic run conditions were 200 V for 40 min. The stain-free gel was then activated and imaged on the Bio-Rad GelDoc Go for total protein loading control. Proteins were wet-transferred from the gel into polyvinylidene difluoride (PVDF) membranes at 30 V overnight. For immunoblotting, DyLight 800-conjugated HA Epitope Tag monoclonal antibody (Thermo Fisher Scientific) was used at dilution 1:5000. The membrane was visualized and imaged with an Odyssey CLx Imager (LICORbio). Images brightness and contrast were adjusted using Fiji ImageJ.

### Statistical analysis

In growth curve experiments, growth rate values (b) were calculated by fitting the growth curve in the exponential phase (from 5 to 15 hours) to an exponential bacterial growth model (OD_600 nm_ ∼ a * 2 ^(b * time)^). Growth rates (growth curve experiments) and relative mRNA levels (RT-qPCR experiments) differences between the treatment and control groups were assessed using a two-tailed Welch’s t-test (n = 3-4 per group). For protoporphyrin IX curve experiments, fluorescence measures (RFU) were normalized by OD_600 nm_. The normalized fluorescence values were transformed to log_2_ scale and differences between the treatment and control groups were assessed using a two-tailed Welch’s t-test (n = 4 per group). Cell area measurements from microscopy images of treatment and control groups were compared using a two-tailed Welch’s t-test (n > 500 per group). For whole RNA-sequencing experiments, pairwise differential expression analysis was performed using Bioconductor package edgeR (3.24.3 with R 3.5.2). Only protein coding genes and long non-coding RNAs (lncRNAs) were considered, and only genes with counts per million values ≥ 0.5 in at least 2 sample were kept for further analyses. Statistical significance was determined by log fold change cutoff of 0.4 and false discovery rate (FDR) cutoff of 0.05. Unless otherwise noted, significance symbols *ns, *, \*\** and *\*\*\** indicate, respectively, P-values > 0.05, < 0.05, < 0.01 and < 0.001 in the corresponding statistical tests.

## Author contributions

F.J.G.G. designed research, performed experiments, analyzed data and wrote the paper; S.S. designed research and wrote the paper.

