## Supporting Information for "A CRISPR Interference System for Inducible Gene Knockdown in soil bacterium *Sinorhizobium meliloti*"

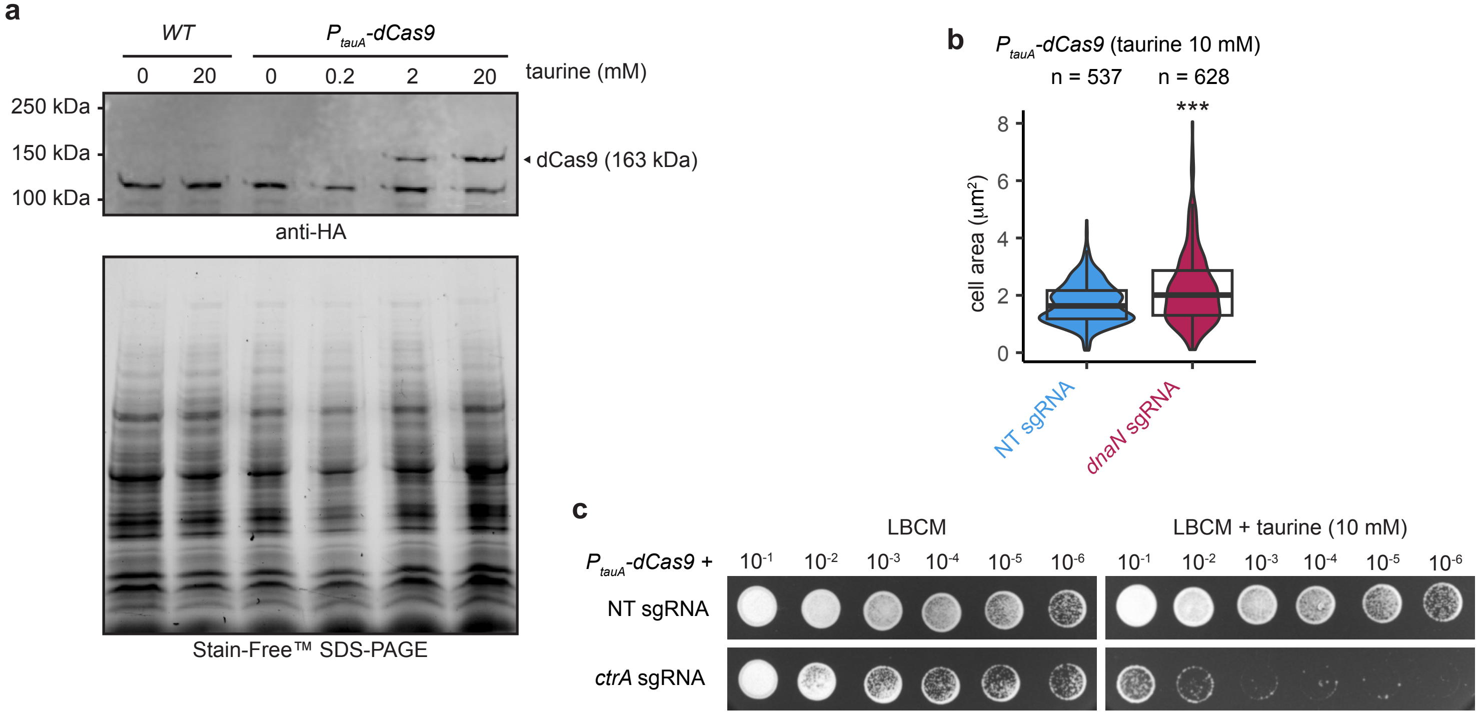


**Fig. S1. a.** *Sth3-dCas9* protein expression under different concentrations of the inducer, taurine. Total protein was separated in an SDS-PAGE Stain-Free™ gel for loading control and HA-tagged *dCas9* was detected using anti-HA antibody through western blot. Expected band size is 163 kDa. **b.** Cell size quantification of *dnaN* knock-down cells from microscopy images. **c.** Viability of strains expressing *PtauA-dCas9* plus non-targeting (NT) or *ctrA* targeting single-guide RNAs (sgRNAs). Shown are serial dilutions of each *S. meliloti* strain on LB media supplemented with $CaCl_2$ and $MgSO_4$ (LBCM) after 3 days at 30°C.

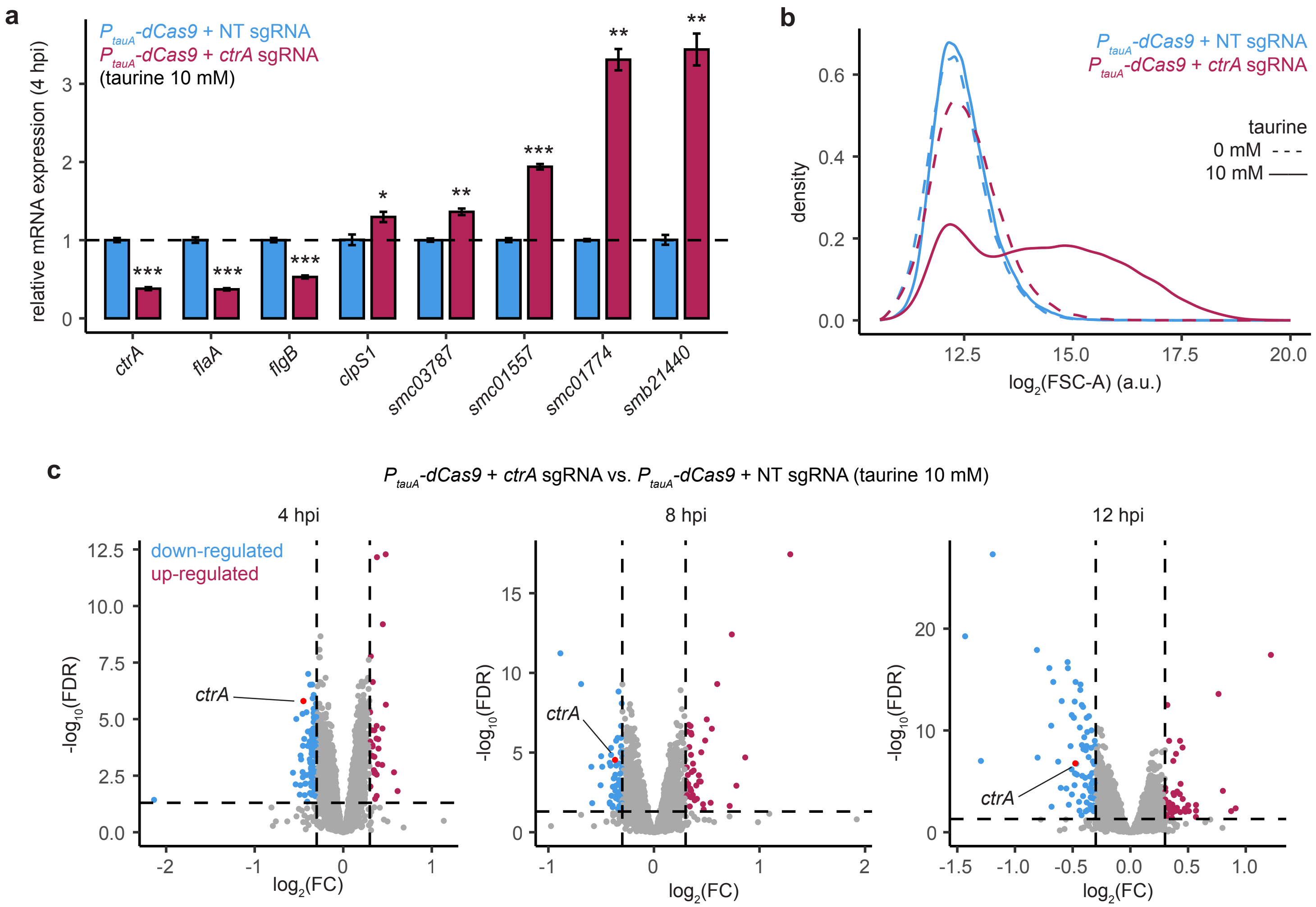


**Fig. S2. a.** Validation of up- and down-regulation of genes selected from *ctrA* knockdown RNA-sequencing experiment. mRNA levels were measured by RT-qPCR at 4 hpi, normalized to *smc00128* levels. **b.** Flow cytometry forward scatter profiles in $\log_2$ scale showing the increased size of *ctrA* knockdown cells at 24 hours after induction of the CRISPRi system. **c.** Volcano plots showing genes significantly up- and down-regulated upon *ctrA* knockdown 4, 8 and 12 hours post induction (hpi), obtained from whole RNA-sequencing. The gene expression fold change between *ctrA* and non-targeting (NT) sgRNA expressing $P_{tauA}$-*dCas9* cells is represented in $\log_2$ scale in the X-axis. Significance is represented in the Y-axis as the $-\log_{10}$ of the p-values. Significance cutoffs: FDR < 0.05, $|\log_2 FC| > 0.4$.

### Supporting Information

### Table S1. List of sgRNA sequences used in this work.

| sgRNA sequence | PAM sequence | Gene ID | Old locus tag | Gene name | Target position (relative to start codon) | Target strand |
|---|---|---|---|---|---|---|
| ACGTCTTTGGCGGGAGTCCGC | - | - | - | - | - | non-targeting (NT) |
| ATCCTTCTAAAAAGAAATAC | CGGCG | *SM_RS14350* | *SMc04019* | *hemH* | -70 | coding strand |
| CGCGGGCGAGGGGACCGCCG | CGGCG | SM_RS01720 | SMc00415 | dnaN | -37 | coding strand |
| AATTCTGATTCACTCTGGCA | AGGTG | SM_RS13920 | SMc00654 | ctrA | -108 | coding strand |
| GAAACGACGCAAGCGACTCG | GGGAG | SM_RS23265 | SMb21440 | - | -77 | coding strand |
| TCGAAACGACTGTCTTCTTA | AGGCG | SM_RS06395 | SMc01906 | hupB | -32 | coding strand |
| TCCTTGTTTTTCTGGCCGAA | GGGCG | SM_RS06395 | SMc01906 | hupB | -169 | coding strand |
| GCCAGAAAAACAAGGAAGAC | CGGCG | SM_RS06395 | SMc01906 | hupB | -165 | template strand |
| CCCTTGCATTCCAAGGGATT | AGGCG | SM_RS06395 | SMc01906 | hupB | -103 | template strand |
| TCCCCTGAACGAGTTGCGAT | GGGAG | SM_RS26970 | SMa0815 | nifA | -43 | coding strand |

### Table S2. List of primers used in this work.

| Primer ID | Name | Sequence |
|---|---|---|
| fgo002 | dCas9_rv | AAGATCGGGTACTTCGAGTC |
| fgo004 | sgRNA_rv | GTCGGCACTTGGTAGCG |
| fgo006 | hemH_sgRNA_2 | ATCCTTCTAAAAAGAAATACGTTTTAGAGCTGTGAAAACAGC |
| fgo014 | dnaN_sgRNA_2 | CGCGGGCGAGGGGACCGCCGGTTTTAGAGCTGTGAAAACAGC |
| fgo025 | tauR_genome_fw | TTGACAAGCTGGATGTCGTC |
| fgo026 | Sth3-dCas9_qPCR_fw | CGAGGTGTTCAAGGACGACA |
| fgo027 | Sth3-dCas9_qPCR_rv | GCCAGCAGCTTCTTCAGGTA |
| fgo028 | Smc00128_qPCR_fw | GATGAAGGACCAGGAAGGTTAC |
| fgo029 | Smc00128_qPCR_rv | CTCGACTCCGCAAGGAAAG |
| fgo039 | gib_fgp003_fw | ACCCAAGTACCGCCACCTAAGCGGGACTCTGGGGTTCGAA |
| fgo040 | gib_fgp003_rv | GTGTGCGTCACCCGGCAACCGTAAGCCCACTGCAAGCTACC |
| fgo041 | gib_GmR_fw | GTAGCTTGCAGTGGGCTTACGGTTGCCGGGTGACGCACAC |
| fgo042 | gib_GmR_rv | TTCGAACCCCAGAGTCCCGCTTAGGTGGCGGTACTTGGGTCG |
| fgo044 | ctrA_sgRNA_2 | AATTCTGATTCACTCTGGCAGTTTTAGAGCTGTGAAAACAGC |
| fgo045 | ctrA_qPCR_fw | TGCGTCTGTCGAAGGTCAAA |
| fgo046 | ctrA_qPCR_rv | CTCGTCCTTGTGGAAAGGCT |
| fgo051 | nifA_sgRNA_1 | TCCCCTGAACGAGTTGCGATGTTTTAGAGCTGTGAAAACAGC |
| fgo055 | dnaN_qPCR_fw | CGTCAGCGAACTGCAGAAAC |
| fgo056 | dnaN_qPCR_rv | GTCATGACGATGGAGCCGAT |
| fgo059 | nifA_qPCR_fw | AGGCCGCAATAGACCAAGTC |
| fgo060 | nifA_qPCR_rv | GATGAAGGCAGTCGGACCAA |
| fgo065 | dual_plasmid_dwn_F | CCAATTCGCCCTATAGTGAGTCGTATTACG |

| | | |
|---|---|---|
| fgo066 | dual_plasmid_up_R | AGAACTAGTGGATCCCCCGGGCTGCAGGAA |
| fgo067 | dual_insert_up_F | TTCCTGCAGCCCGGGGGATCCACTAGTTCTCAAATAAAACGAAAGGCTCAGTCGAAAGAC |
| fgo068 | dual_insert_short_dwn_R | CGTAATACGACTCACTATAGGGCGAATTGGAAGACCCGTTTATAAAACGAAAGGCTCAGT |
| fgo078 | hupB_sgRNA_1 | TCGAAACGACTGTCTTCTTAGTTTTAGAGCTGTGAAAACAGC |
| fgo079 | hupB_sgRNA_2 | TCCTTGTTTTTCTGGCCGAAGTTTTAGAGCTGTGAAAACAGC |
| fgo080 | hupB_sgRNA_3 | GCCAGAAAAACAAGGAAGACGTTTTAGAGCTGTGAAAACAGC |
| fgo081 | hupB_sgRNA_4 | CCCTTGCATTCCAAGGGATTGTTTTAGAGCTGTGAAAACAGC |
| fgo085 | nifB_qPCR_fw | GGCATCCTCACCAAGGTCAA |
| fgo086 | nifB_qPCR_rv | TCCCGTGTATGCGGTTTGAA |
| fgo087 | nifT_qPCR_fw | TCGAGGAGCCGATCATTTCG |
| fgo088 | nifT_qPCR_rv | GCCGGATATTTGCCTCGACT |
| fgo099 | hupB_qPCR_fw | TCTTCTGCCGTTGATGCAGT |
| fgo100 | hupB_qPCR_rv | GAGAAATTGCCGAAGCCGAC |
| fgo101 | SMb21440_qPCR_fw | TCCACGAAAATCGGGGCTAC |
| fgo102 | SMb21440_qPCR_rv | CCGCCTTGACCATTTCCTGA |
| fgo103 | flaA_qPCR_fw | CCTTCGACGGAGACTATGCC |
| fgo104 | flaA_qPCR_rv | TAGACGACTTCCTGACCCGT |
| fgo105 | flgB_qPCR_fw | AACATCGCCAATGCCAACAC |
| fgo106 | flgB_qPCR_rv | ATCTGGATGCCGGTGTTCTG |
| fgo107 | SMc01557_qPCR_fw | TATGTTGCGGATGTGACGGG |
| fgo108 | SMc01557_qPCR_rv | TTGGCGAAAACGATCGCCTG |
| fgo111 | SMc03787_qPCR_fw | CGGCTGTTTTCCCAATGTCG |
| fgo112 | SMc03787_qPCR_fw | GTCTGAGGCGGGTCATGAAA |
| fgo113 | clpS1_qPCR_fw | CGAAGACCAAGAAGCCGAGT |
| fgo114 | clpS1_qPCR_rv | CTGAAAGAAGCGCTCGAGGA |
| fgo117 | SMc01774_qPCR_fw | GCGAAAATCTCCGTCATCGC |
| fgo118 | SMc01774_qPCR_rv | GCCAGTTCATTTCGACGCTC |
| fgo119 | SMb21440_sgRNA_1 | GAAACGACGCAAGCGACTCGGTTTTAGAGCTGTGAAAACAGC |

### Table S3. List of plasmids used in this work.

| Plasmid ID | Resistance | Expression | Origin |
|---|---|---|---|
| fgp003 | Kan/Neo | NT-sgRNA | Guzzo et al. 2020, Addgene #133339 |
| fgp004 | Gm | NT-sgRNA | fgp003 |
| fgp005 | Kan/Neo | $P_{tauA}$-Sth3-dCas9-HA | Mostofavi et al. 2014,<br>Guzzo et al. 2020 |
| fgp009 | Gm | hemH-sgRNA_2 | fgp004 |
| fgp011 | Gm | ctrA-sgRNA_2 | fgp004 |
| fgp016 | Gm | dnaN-sgRNA_2 | fgp004 |
| fgp023 | Gm | nifA-sgRNA_1 | fgp004 |

| | | | |
|---|---|---|---|
| fgp026 | Gm | hemH-sgRNA_2<br>ctrA-sgRNA_2 | fgp009, fgp011 |
| fgp031 | Gm | hupB-sgRNA_1 | fgp004 |
| fgp032 | Gm | hupB-sgRNA_2 | fgp004 |
| fgp033 | Gm | hupB-sgRNA_3 | fgp004 |
| fgp034 | Gm | hupB-sgRNA_4 | fgp004 |
| fgp038 | Gm | hupB-sgRNA_1<br>hupB -sgRNA_4 | fgp031, fgp034 |
| fgp039 | Gm | hupB-sgRNA_1<br>hupB-sgRNA_2<br>hupB -sgRNA_4 | fgp031, fgp032, fgp034 |
| fgp042 | Gm | Smb21440-sgRNA_1 | fgp004 |
| fgp045 | Gm | ctrA-sgRNA_2<br>Smb21440-sgRNA_1 | fgp011, fgp042 |

### Table S4. List of strains used in this work.

| Strain ID | Organism | Origin | Resistance | Expression |
|---|---|---|---|---|
| fgs001 | *Sinorhizobium meliloti* 1021 | Meade et al. 1982 | Strep | - |
| fgs004 | *Sinorhizobium meliloti* 1021 | fgs001 | Strep, Neo | $P_{tauA}$-dCas9 |
| fgs006 | *Escherichia coli* | - | Clo, Kan | pRK600 (helper strain) |
| fgs009 | *Sinorhizobium meliloti* 1021 | 1021 | Strep, Gen | NT-sgRNA |
| fgs014 | *Sinorhizobium meliloti* 1021 | fgs004 | Strep, Neo, Gen | $P_{tauA}$-dCas9 + NT-sgRNA |
| fgs016 | *Sinorhizobium meliloti* 1021 | fgs004 | Strep, Neo, Gen | $P_{tauA}$-dCas9 + hemH-sgRNA-2 |
| fgs018 | *Sinorhizobium meliloti* 1021 | fgs004 | Strep, Neo, Gen | $P_{tauA}$-dCas9 + ctrA-sgRNA-2 |
| fgs021 | *Sinorhizobium meliloti* 1021 | fgs004 | Strep, Neo, Gen | $P_{tauA}$-dCas9 + dnaN-sgRNA-2 |
| fgs047 | *Sinorhizobium meliloti* 1021 | fgs004 | Strep, Neo, Gen | $P_{tauA}$-dCas9 + nifA-sgRNA-1 |
| fgs053 | *Sinorhizobium meliloti* 1021 | fgs004 | Strep, Neo, Gen | $P_{tauA}$-dCas9 + hemH-sgRNA-2 + ctrA-sgRNA-2 |
| fgs058 | *Sinorhizobium meliloti* 1021 | fgs004 | Strep, Neo, Gen | $P_{tauA}$-dCas9 + hupB-sgRNA-1 |
| fgs059 | *Sinorhizobium meliloti* 1021 | fgs004 | Strep, Neo, Gen | $P_{tauA}$-dCas9 + hupB-sgRNA-2 |
| fgs060 | *Sinorhizobium meliloti* 1021 | fgs004 | Strep, Neo, Gen | $P_{tauA}$-dCas9 + hupB-sgRNA-3 |
| fgs061 | *Sinorhizobium meliloti* 1021 | fgs004 | Strep, Neo, Gen | $P_{tauA}$-dCas9 + hupB-sgRNA-4 |
| fgs062 | *Sinorhizobium meliloti* 1021 | fgs004 | Strep, Neo, Gen | $P_{tauA}$-dCas9 + hupB-sgRNA-1 + hupB-sgRNA-4 |
| fgs063 | *Sinorhizobium meliloti* 1021 | fgs004 | Strep, Neo, Gen | $P_{tauA}$-dCas9 + hupB-sgRNA-1 + hupB-sgRNA-2 + hupB-sgRNA-4 |
| fgs067 | *Sinorhizobium meliloti* 1021 | fgs004 | Strep, Neo, Gen | $P_{tauA}$-dCas9 + Smb21440-sgRNA-1 |
| fgs073 | *Sinorhizobium meliloti* 1021 | fgs004 | Strep, Neo, Gen | $P_{tauA}$-dCas9 + ctrA-sgRNA-2 + Smb21440-sgRNA-1 |
